# Phenotypic Characterization of Signature-Tagged Mutants Identifies Physiological Determinants of *Vibrio vulnificus* Fitness

**DOI:** 10.64898/2025.12.18.695119

**Authors:** Kohei Yamazaki, Takehiro Kado, Takashige Kashimoto

**Author notes:** Address correspondence to Takashige Kashimoto. Kohei Yamazaki and Takehiro Kado contributed equally to this work. Author order was determined in order of increasing seniority.

## Abstract

*Vibrio vulnificus* is an opportunistic marine pathogen that causes severe wound-associated and systemic infections. Following entry into the host, the bacterium must rapidly adapt to host-associated stresses that differ substantially from those encountered in aquatic environments. However, the physiological functions supporting bacterial fitness during infection remain incompletely understood. Previously, we applied signature-tagged mutagenesis (STM) to identify genes required for *V. vulnificus* survival during host infection, revealing numerous loci that did not correspond to classical toxin-encoding genes. In the present study, we extended this genome-wide screen by linking STM-identified mutations to observable fitness-related phenotypes. Functional annotation revealed enrichment of genes associated with chemotaxis, flagellar motility, regulation, metabolism, and poorly characterized functions. Phenotypic analyses showed that many STM-derived mutants exhibited defects in swimming motility and altered colony surface properties. Bioluminescence imaging further revealed distinct patterns of impaired persistence and dissemination within host tissues, while several mutants displayed increased susceptibility to phagocytic stress in an HL-60-derived neutrophil model. Notably, some regulatory mutants affecting global signaling pathways exhibited impaired tissue dissemination despite retaining resistance to phagocytic stress, indicating the presence of fitness determinants that operate independently of classical surface-associated or cytotoxic traits. Together, these findings demonstrate that *V. vulnificus* fitness during infection depends on diverse physiological pathways beyond classical virulence factors, highlighting the value of phenotype-centered analyses for understanding bacterial adaptation in host-associated environments.

**Importance:** *Vibrio vulnificus* causes rapidly progressive wound infections and septicemia, yet the bacterial functions that support fitness within host environments remain incompletely defined. While substantial effort has focused on canonical virulence factors and regulation, increasing evidence suggests that successful infection also depends on broader physiological adaptation. In this study, we link signature-tagged mutagenesis with systematic phenotypic analyses to define physiological determinants of *V. vulnificus* fitness during host-associated infection. Our results demonstrate that genes involved in motility, regulatory signaling, metabolism, and stress tolerance collectively shape bacterial persistence and dissemination in host tissues. Notably, we identify regulatory and metabolic determinants that influence fitness independently of classical surface-associated or cytotoxic traits, highlighting noncanonical pathways that contribute to pathogenic success. By integrating genome-wide screening with phenotype-centered analyses, this work advances understanding of how physiological adaptation underpins *V. vulnificus* infection and provides a framework for studying bacterial fitness alongside established concepts of bacterial virulence.

## Introduction

The public health relevance of *Vibrio vulnificus* has increased markedly in recent years. Rising sea surface temperatures associated with global climate change have expanded the geographical range of this marine bacterium, leading to increased case numbers and reports of severe wound-associated infections in regions that were previously unaffected (1, 2). Because *V. vulnificus* infection is frequently associated with rapidly progressive necrotizing soft tissue infections and high mortality, particularly following wound exposure, it represents a growing public health concern worldwide (3–8). As opportunities for environmental exposure continue to grow, there is an increasing need to understand how *V. vulnificus* adapts to host-associated environments following entry into the human body. *Vibrio vulnificus* is a marine bacterium that opportunistically invades host tissues through wounds, where it encounters physicochemical constraints, nutrient limitation, and host-derived biotic pressures that differ substantially from those in external aquatic environments. Successful proliferation in soft tissues is known to depend on several bacterial traits that support survival under these stresses. The RTX toxin has been widely reported to exert cytotoxic effects on host immune cells, particularly neutrophils, thereby impairing early innate immune defenses (9, 10). In addition, the capsular polysaccharide contributes to resistance against phagocytosis, promoting bacterial persistence in host environments (11).

Chemotaxis and flagellar motility further enable bacterial migration and spatial expansion within infected tissues, facilitating access to niches permissive for growth (12).

Although these traits are established contributors to pathogenicity, they represent only a subset of the bacterial functions required for survival in complex host-associated environments. Increasing evidence from genome-wide analyses indicates that many genes essential for *in vivo* fitness do not correspond to classical virulence determinants but instead encode physiological functions involved in regulation, motility, and metabolism ( 13–16). These findings suggest that environmental adaptation, rather than toxin-mediated damage alone, plays a central role during early stages of infection.

Previously, we applied signature-tagged mutagenesis (STM) to identify genes required for *V. vulnificus* fitness during wound infection (13). That study generated a genome-wide map of candidate loci contributing to survival within host tissues and revealed that many STM-identified genes were not canonical virulence factors. Instead, a large proportion of the recovered loci encoded components of motility systems, regulatory elements, metabolic enzymes, or proteins of unknown function, supporting the idea that physiological adaptation is critical for persistence *in vivo*.

In the present study, we extend this work by focusing on phenotypic characterization of representative STM-identified mutants. To directly link STM selection with observable fitness-related traits, we employed *in vivo* bioluminescence imaging to monitor bacterial persistence and dissemination in real time, together with assays measuring tolerance to phagocyte-associated stress.

This approach allows evaluation of how specific physiological pathways contribute to environmental fitness in host tissues without addressing the molecular mechanisms underlying individual virulence factors.

Using this strategy, we show that mutants defective in motility ( *motX*), regulatory signaling (*barA* and *luxO*), and amino-sugar metabolism (*gpsK*) exhibit reduced *in vivo* fitness and diminished tolerance to phagocytic stress. Together, these findings update and extend our previous STM analysis by integrating functional phenotyping and real-time imaging, providing insight into the physiological systems that support *V. vulnificus* adaptation to host-associated environments.

## Methods

### Animals and Ethics Statement

Five-week-old female C57BL/6 or BALB/c mice (Charles River Laboratories Japan) were housed under SPF conditions with ad libitum access to food and water, following a 12:12 h light-dark cycle. All animal experiments were approved by the Institutional Animal Care and Use Committee of Kitasato University, in accordance with JALAS guidelines, consistent with our previous reports (Approval No. 19-218).

### Bacterial Strains

*Vibrio vulnificus* CMCP6 (clinical isolate) was used as the wild-type (WT) strain. Transposon mutants were generated as described previously (13). Briefly, *Escherichia coli* BW19795 carrying the signature-tagged mini-Tn5Km2 transposon on the suicide vector pUT was used as the donor strain for conjugation. Transposons were introduced into *Vibrio vulnificus* by biparental mating, and transconjugants were selected on LB agar plates containing kanamycin under appropriate conditions. For genetic complementation, the target gene was amplified by PCR and cloned into the low-copy-number plasmid pACYC184 using In-Fusion cloning reactions (Clontech, TaKaRa, Shiga, Japan), according to the manufacturer’s instructions. The resulting complementing plasmid was introduced into *V. vulnificus* by electroporation. Transformants were selected on LB agar plates containing chloramphenicol (10 µg/ml) and incubated overnight at 37°C. All strains were grown aerobically in Luria–Bertani (LB) broth or on LB agar at 37 °C, as described previously. When appropriate, antibiotics were added at the following concentrations: rifampicin 50 µg/ml, kanamycin 100 µg/ml, or ampicillin 100 µg/ml.

### Growth and Inoculation Conditions

Overnight cultures were diluted 1:20 into fresh LB medium and incubated for 2 h at 37°C with agitation (163 rpm). Bacterial cells were harvested, washed with PBS containing 0.1% gelatin, and resuspended to the desired concentration. Mice were inoculated subcutaneously in the caudal thigh with 1 × 10⁶ CFU of wild-type or mutant strains. Infected mice were monitored at defined time points (3 –12 h postinfection) for clinical signs.

### Identification of transposon insertion sites

The transposon insertion sites were determined using arbitrarily primed PCR as described previously (13). Briefly, genomic DNA was isolated from each transposon mutant, and PCR was performed using primers specific to the transposon together with arbitrary primers targeting the *Vibrio vulnificus* genome. The resulting PCR products were sequenced, and insertion sites were identified by sequence homology searches against the *V. vulnificus* genome database.

### Motility assay

Swimming motility was assessed on 0.3% agar, representing a low-viscosity environment relevant to tissue and fluid interfaces (12,13).

### Capsule Opacity Assay (CPS phenotype)

Overnight cultures were spotted onto LB agar and incubated 12 h at 37 °C. Colony translucency/opacity was visually inspected (11,13).

### *In vivo* bioluminescence imaging

To evaluate persistence and dissemination, WT and mutants were transformed with pXen-13 (*luxCDABE*). Bioluminescent WT and mutant strains were visualized using an IVIS-200 imaging system (PerkinElmer). Mice were anesthetized with isoflurane, and images were acquired with a fixed exposure time of 1 minute at 3, 6, 9, and 12 hours after subcutaneous infection (15,17).

### Opsonophagocytic survival assay

Neutrophil-like cells were generated by differentiating HL-60 cells with 1.25% dimethyl sulfoxide (DMSO) for 5 days, as described previously (16). Bacterial cells were incubated with differentiated HL-60 cells at a multiplicity of infection (MOI) of 1:20 in the presence of 10% human serum for 45 min at 37°C with gentle rotation. Following incubation, host cells were lysed by treatment with 0.05% saponin, and samples were serially diluted and plated onto LB agar for enumeration of surviving CFU. Bacterial survival was calculated by comparing CFU recovered after incubation with HL-60 cells to CFU recovered from control samples incubated for the same duration under identical conditions in the absence of HL-60 cells.

### Statistical analysis

All statistical analyses were performed using GraphPad Prism (version 8). Quantitative comparisons of bacterial CFU counts were evaluated using the Mann–Whitney U test, as the data did not follow a normal distribution. For all experiments, *p* values of < 0.05 were considered statistically significant.

## Results

### Functional categorization of STM-identified genes

To evaluate the functional characteristics of genes identified by the STM screen, we first examined the distribution of annotated functions among the selected loci. All STM-derived mutants exhibited growth kinetics comparable to WT under standard in vitro culture conditions (data not shown), indicating that the observed attenuation was not attributable to intrinsic growth defects. The STM-attenuated gene set was enriched for genes associated with chemotaxis and flagellar motility, including multiple components of the basal body, motor complex, hook, and chemotactic signaling pathways (Table 1). In addition to motility-associated genes, the STM-identified loci included regulatory elements, metabolic enzymes, stress response proteins, and factors involved in chromosome maintenance and cell division (Table 1). A substantial fraction of the identified genes encoded proteins annotated as hypothetical or with limited functional characterization.

**Table 1.**
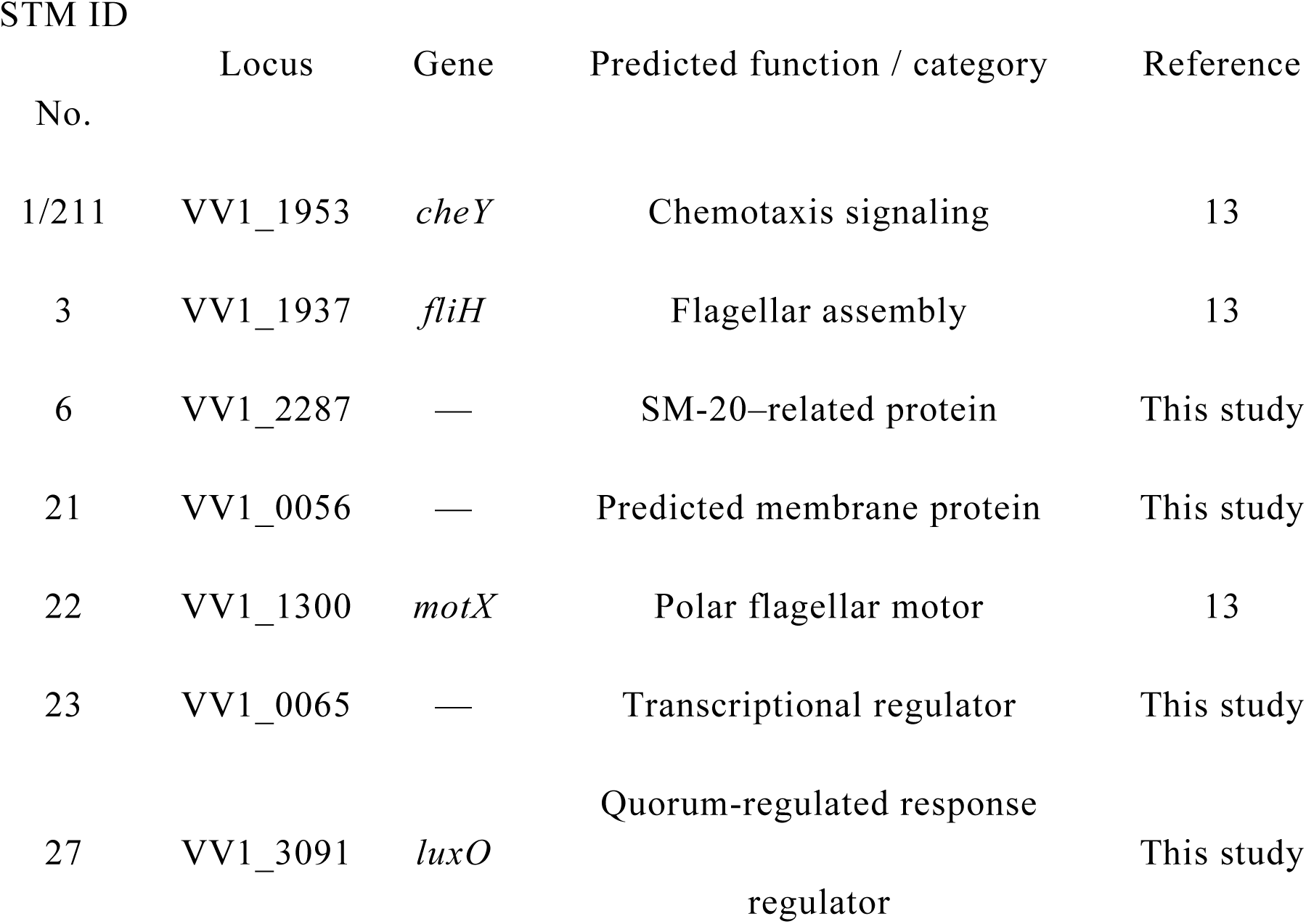
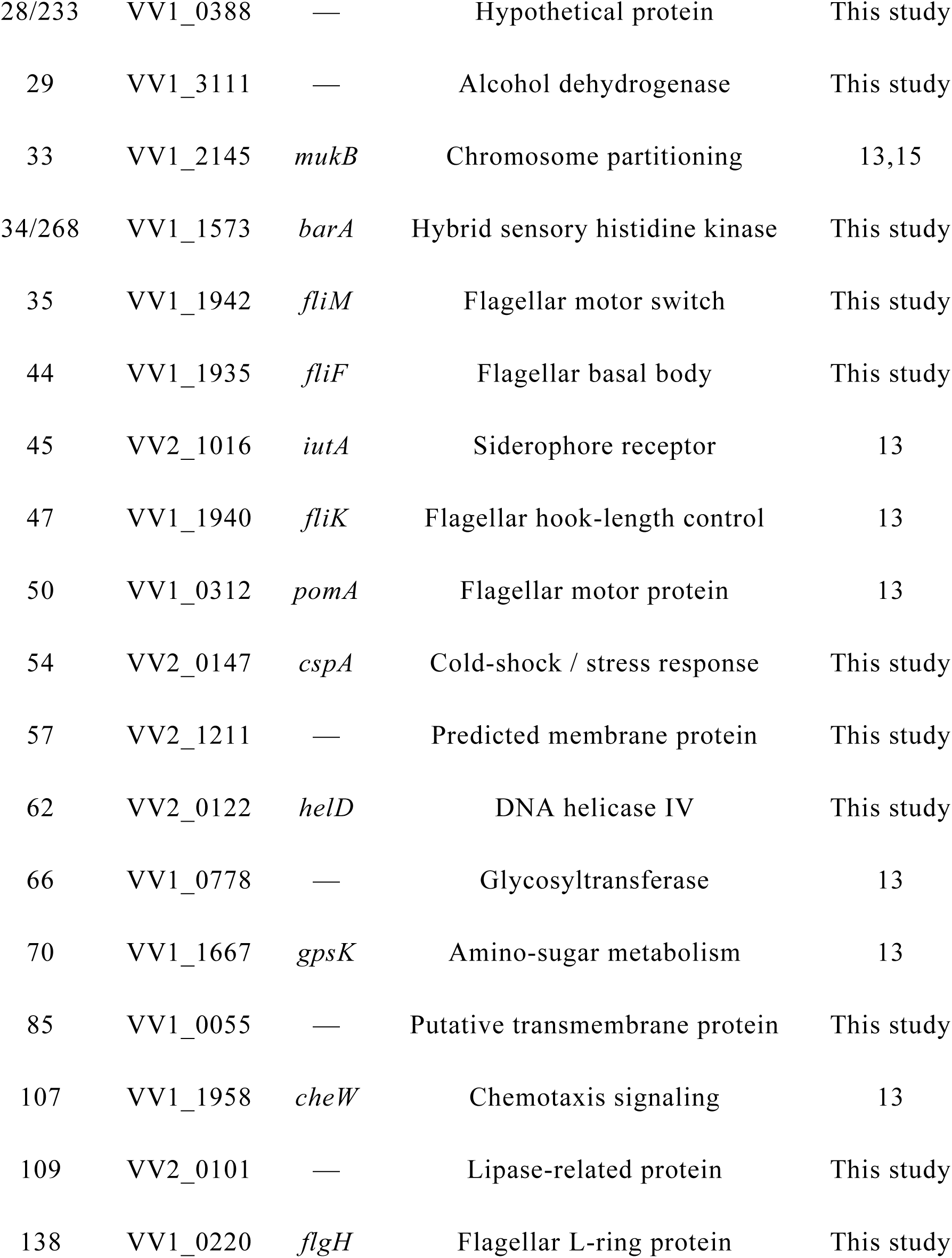
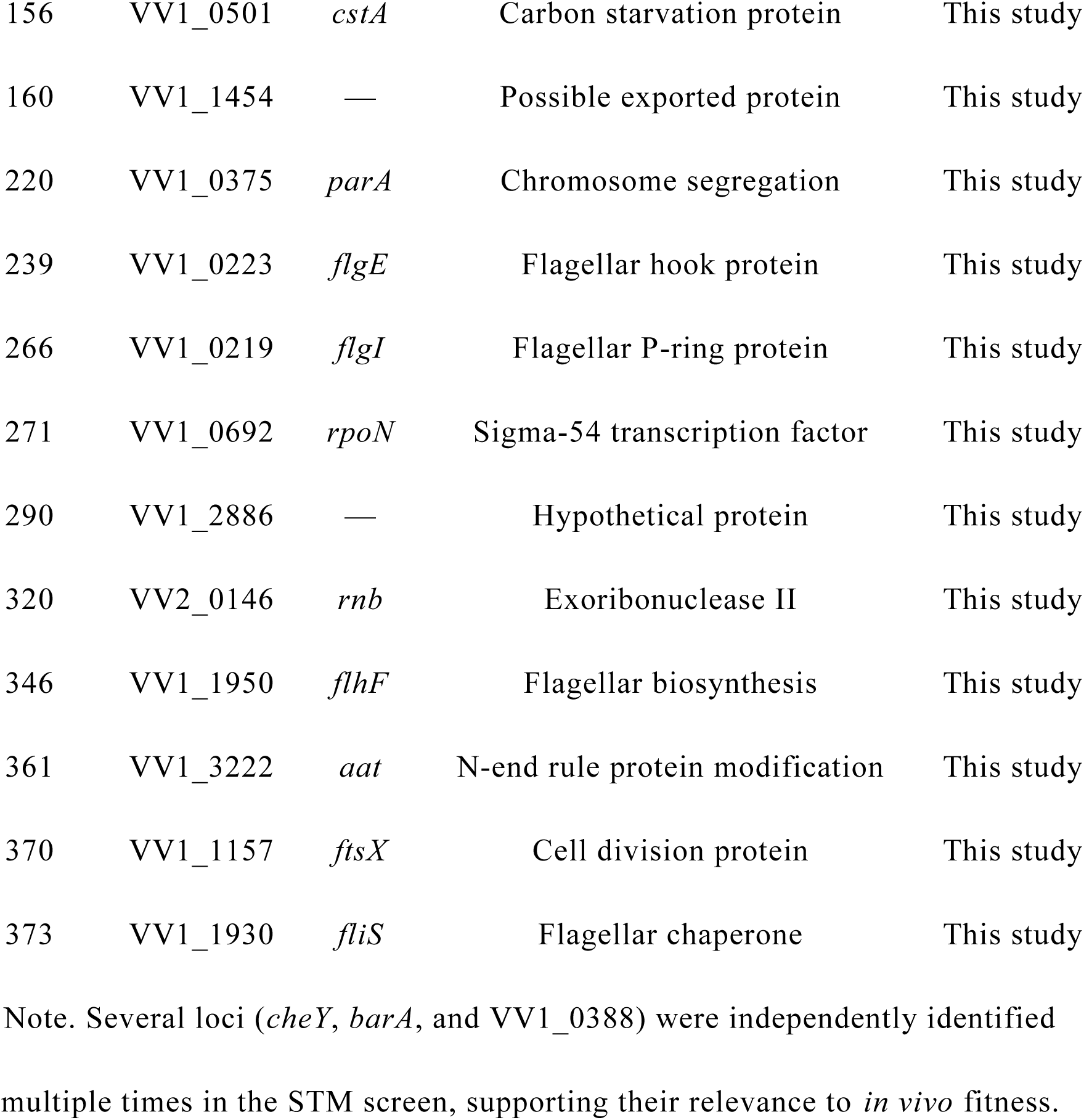
STM-identified genes selected for phenotypic characterization.

### Motility defects among STM-identified mutants

To assess whether STM-identified mutations were associated with defects in motility-related phenotypes, all mutants listed in Table 1 were subjected to a swimming motility assay as an initial phenotypic screen. WT exhibited robust radial expansion on soft-agar plates, whereas multiple STM-derived mutants displayed marked reductions in motility (Fig. 1).

**Figure 1.**
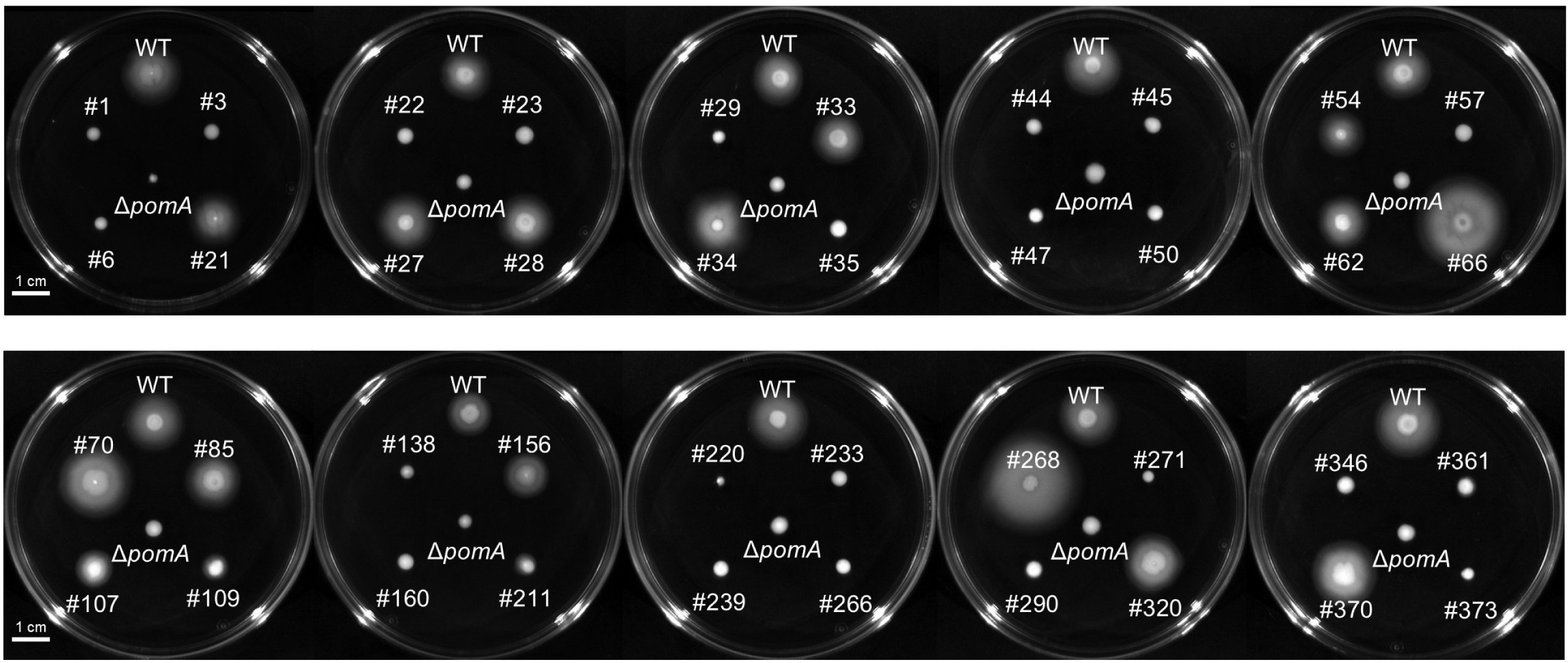
Swimming motility of STM-derived mutants. Swimming motility of WT strain and STM-derived mutants assessed on soft-agar plates. Images show radial expansion following incubation under identical conditions.

As expected, disruptions in genes encoding flagellar structural components and motor-associated proteins, including *motX, fliH, fliM, fliF, fliK, flgE, flgH, flgI,* and *flhF*, resulted in severe motility defects (Fig. 1). Mutations in chemotactic signaling proteins (*cheY* and *cheW*) similarly impaired swimming behavior, consistent with their established roles in directed bacterial movement. In addition to these canonical motility-related genes, several mutants carrying insertions in genes not directly annotated as motility components also exhibited reproducible reductions in swimming motility. These included genes encoding a siderophore receptor (*iutA*), predicted membrane proteins, lipase-related proteins, exported proteins, chromosome segregation factors (*parA*), the sigma-54 transcription factor (*rpoN*), and an N-end rule protein modification enzyme (*aat*) (Fig. 1). These findings indicate that diverse physiological and regulatory functions contribute indirectly to motility-associated behaviors in *V. vulnificus*.

### Capsule-associated colony opacity variations

Colony morphology was assessed as an indicator of surface-associated properties potentially relevant to host adaptation. WT formed opaque colonies on LB agar, whereas several STM-derived mutants produced colonies with increased translucency (Fig. 2). Notably, mutants carrying insertions in the regulatory signaling gene *barA* and a gene encoding a glycosyltransferase exhibited a translucent colony phenotype comparable to that observed in the environmental isolate control strain E4 (Fig. 2). This altered colony morphology was reproducibly observed in mutants affecting regulatory signaling pathways and glycosylation-related functions. In contrast, STM-derived mutants disrupted in flagellar motility or chemotactic signaling genes retained colony opacity comparable to that of the wild-type strain, indicating that the observed translucency phenotype was not a general consequence of STM mutagenesis but was associated with specific functional categories.

**Figure 2.**
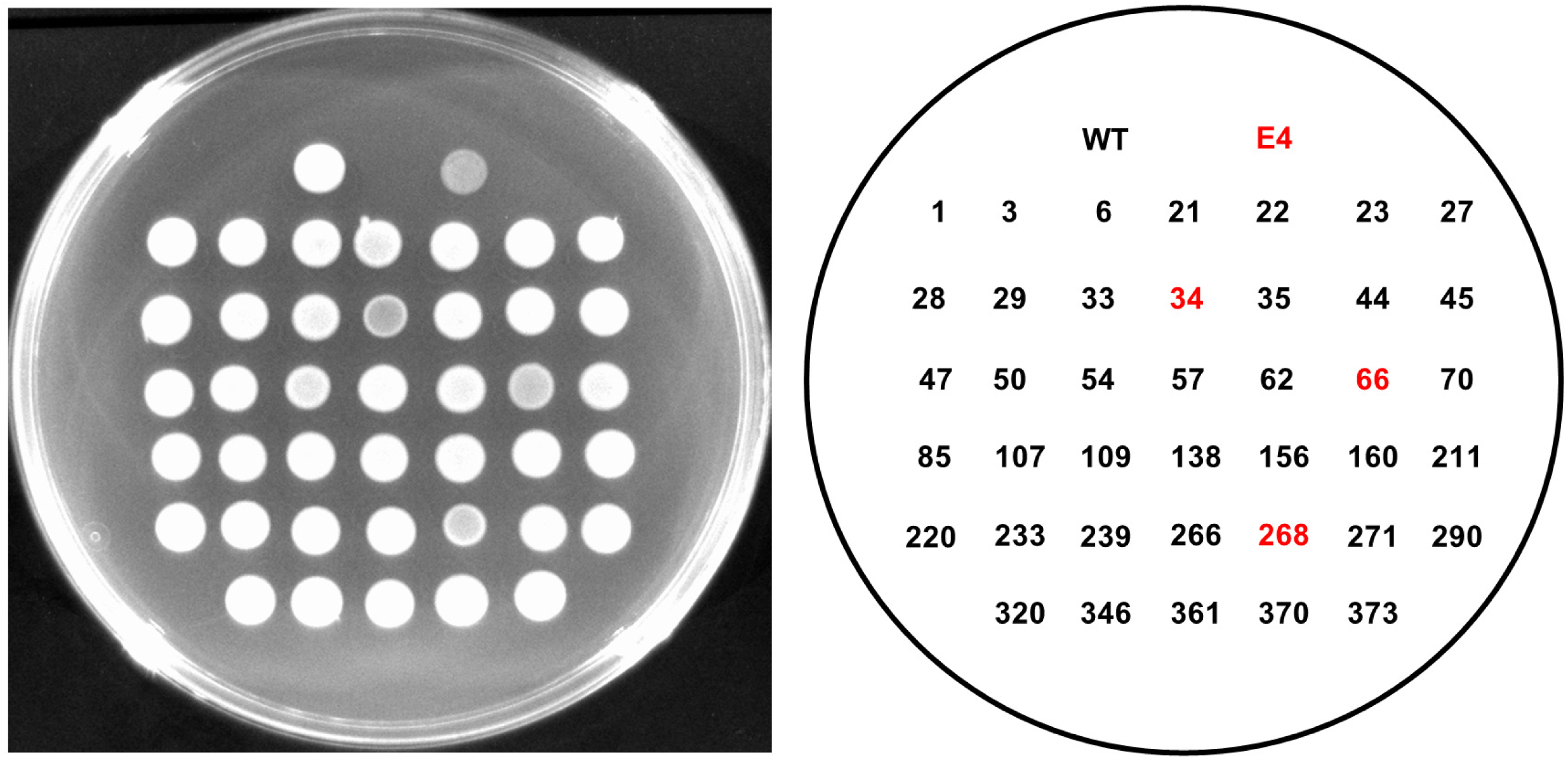
Colony surface properties of STM-derived mutants. Colony morphology of WT and STM-derived mutants grown on LB agar plates. Representative images illustrate differences in colony opacity and surface appearance. The left panel shows colony phenotypes, and the right panel indicates the corresponding mutant identification numbers.

### Reduced persistence and dissemination in soft tissues among representative STM-derived mutants

To determine whether STM-identified mutations impaired bacterial fitness in soft tissues, bioluminescent derivatives of selected mutants were monitored during subcutaneous infection using *in vivo* bioluminescence imaging (IVIS). WT exhibited progressive dissemination of luminescent signals from the inoculation site, followed by signal attenuation by 12 h postinfection, consistent with active persistence, expansion, and subsequent invasion into deeper soft tissues (Fig. 3) (12, 17).

**Figure 3.**
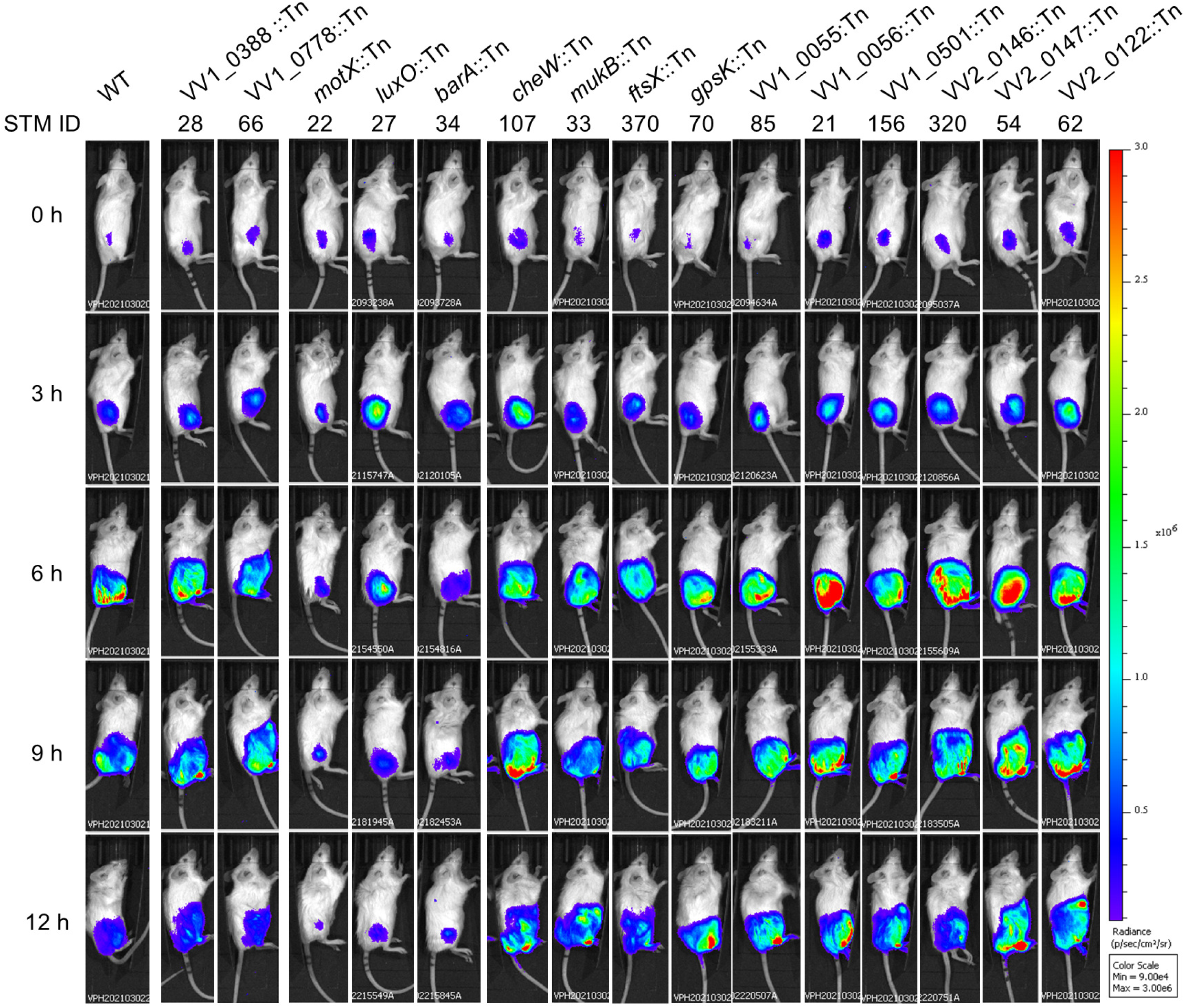
Bioluminescence imaging of STM-derived mutants during subcutaneous infection. Bioluminescent signals of WT and selected STM-derived mutants were monitored at the indicated time points following subcutaneous infection. Images were acquired using identical imaging parameters.

Based on IVIS signal dynamics, STM-derived mutants could be broadly classified into three phenotypic groups. The first group displayed behavior in soft tissues comparable to that of WT. Mutants carrying insertions in VV1_ 0388 and VV1_0778 showed sustained luminescent signals and spatial dissemination similar to those observed for WT, indicating that disruption of these loci did not markedly impair persistence or spread in soft tissues.

The second group exhibited severe defects in dissemination from the infection site. As expected, the motility-deficient mutant *motX::Tn* failed to spread within soft tissues, consistent with previous observations that bacterial motility is essential for expansion and dissemination in this infection model (12). Similarly, mutants disrupted in regulatory signaling pathways ( *luxO::Tn* and *barA::Tn*) showed restricted luminescent signals that remained localized near the inoculation site, resembling the phenotype of the motility-deficient mutant (13).

The third group showed limited dissemination accompanied by prolonged persistence of luminescent signals at the infection site. The chemotaxis-deficient mutant *cheW::Tn* exhibited spread within soft tissues; however, luminescent signals remained detectable over the observation period. This phenotype is consistent with previously described behavior of chemotaxis mutants that retain motility but exhibit impaired invasion into deeper tissue compartments (12). A similar IVIS phenotype was observed for several additional STM-derived mutants, including *mukB*::Tn*, ftsX*::Tn*, gpsK*::Tn, VV1_0055::Tn, VV1_0056::Tn, VV2_0147::Tn, VV2_0122::Tn, VV1_0501::Tn, and VV2_0146::Tn.

Collectively, IVIS analysis revealed distinct phenotypic classes among STM-derived mutants in soft tissues, reflecting differential defects in bacterial persistence, dissemination within soft tissues, and invasion into deeper tissue compartments.

### Increased susceptibility of STM-derived mutants to phagocytic stress

Because early host responses to subcutaneous infection involve rapid recruitment of phagocytic cells, STM-derived mutants were examined for tolerance to phagocytic stress using differentiated HL-60 cells as a neutrophil-like cell model. WT demonstrated substantial survival following exposure to phagocytic cells (Fig. 4). The *barA*::Tn mutant exhibited survival comparable to that of WT (Fig. 4), indicating that disruption of *barA* did not markedly impair resistance to phagocytic stress under these conditions.

**Figure 4.**
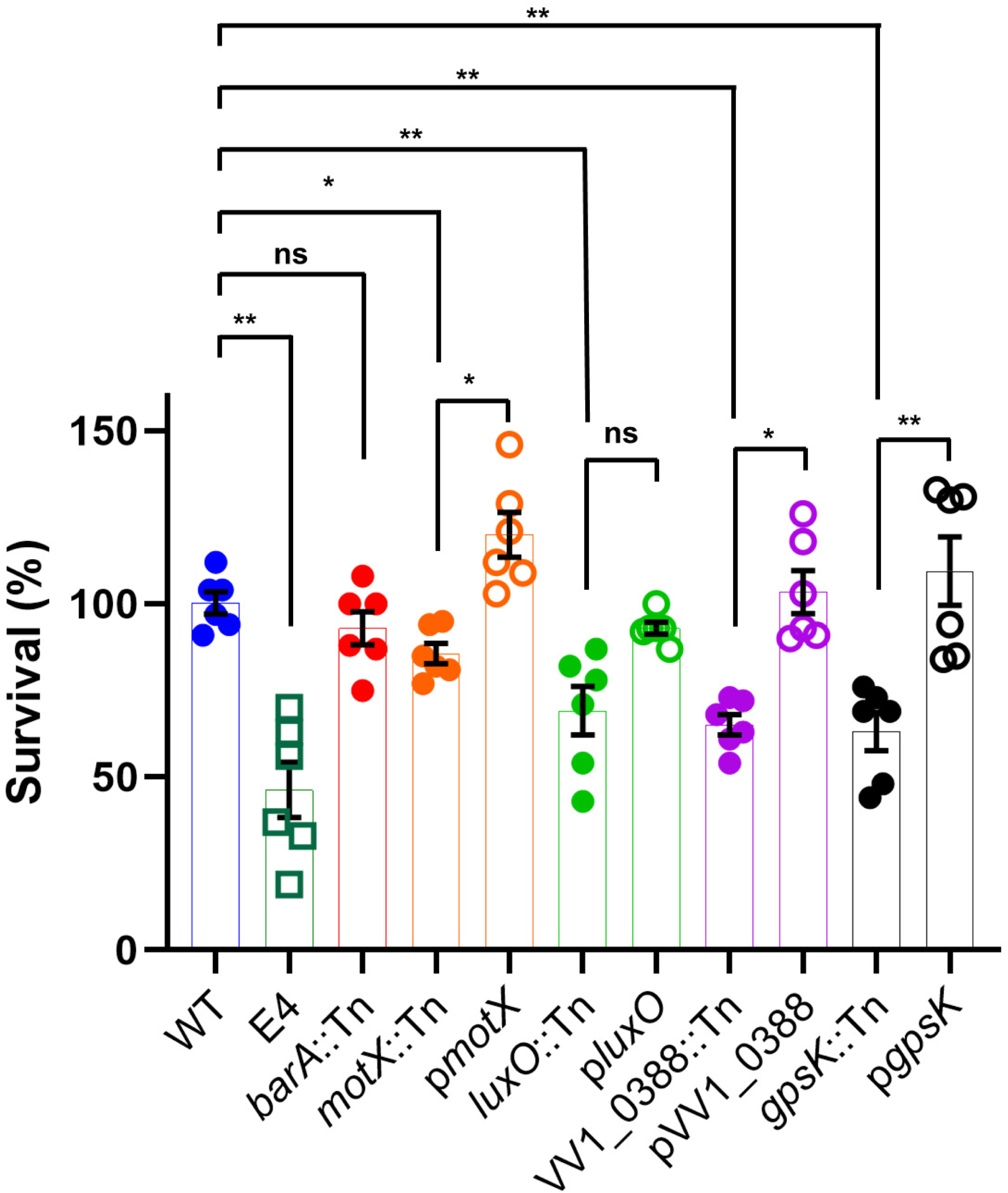
Tolerance of STM-derived mutants to phagocytic stress. Survival of WT and STM-derived mutants following exposure to HL-60-derived neutrophils. Data are presented as percent survival relative to control samples incubated under identical conditions in the absence of HL-60 cells. Statistical significance was evaluated using the Mann–Whitney U test; \**P* < 0.05 and \*\**P* < 0.01.

In contrast, several STM-derived mutants showed significantly reduced survival after incubation with HL-60 cells. Mutants carrying insertions in *motX*, *luxO*, VV1_0388, and *gpsK* displayed pronounced decreases in survival relative to WT (Fig. 4), indicating increased susceptibility to phagocyte-associated stress. These reductions were consistently observed across independent experiments.

To confirm that the observed phenotypes were attributable to the respective transposon insertions, complementation analyses were performed. Introduction of the corresponding wild-type genes restored survival to levels comparable to WT, confirming that the increased susceptibility to phagocytic stress resulted from disruption of the targeted loci.

## Discussion

In this study, we extended our previous STM-based analysis by integrating targeted phenotypic characterization to define physiological traits associated with *Vibrio vulnificus* fitness in host-associated environments. By combining STM selection with motility assays, evaluation of surface-associated properties, bioluminescence-based imaging in soft tissues, and phagocytic stress assays, we directly linked STM-identified loci to observable fitness-related phenotypes without addressing the molecular mechanisms underlying individual virulence factors.

The functional composition of STM-identified genes provides initial validation of the screening strategy. The enrichment of chemotaxis- and flagellar motility–associated genes among STM-attenuated mutants is consistent with the established importance of directed movement for bacterial expansion within soft tissues (Table 1) (12, 13). Motility enables bacteria to explore heterogeneous host environments and access niches permissive for growth, and its repeated recovery in STM-based screens supports the view that motility-related functions represent core physiological requirements for fitness during early infection (12). However, the motility-deficient *motX* mutant retained substantial resistance to phagocytic stress (Fig. 4). In the HL-60 survival assay, *motX*::Tn exhibited a median survival of 83.5%, indicating that the absence of motility alone does not render *V. vulnificus* completely susceptible to phagocytic killing. This residual resistance is likely supported by additional virulence-associated traits retained by the *motX* mutant, such as capsular polysaccharide and RTX toxin production (9,10,12).

In contrast, genes encoding the RTX toxin were not recovered among STM-selected loci (Table 1). This absence likely reflects a methodological characteristic of STM-based negative selection screens rather than a lack of contribution of RTX to virulence. Because STM involves pooled infection with multiple mutant strains, secreted virulence factors such as toxins can be functionally complemented by neighboring bacteria. Under these conditions, mutants defective in toxin production may not exhibit a competitive disadvantage and therefore escape negative selection. Consequently, STM preferentially identifies cell-autonomous physiological functions required for survival and persistence, while diffusible virulence factors are underrepresented (13,14).

Beyond motility-associated genes, the STM-identified set included regulatory elements, metabolic enzymes, stress response proteins, and factors involved in chromosome maintenance and cell division (15,16,18–23). Notably, a substantial fraction of the recovered loci encoded proteins of unknown or poorly characterized function. The recurrence of such genes among STM-attenuated mutants suggests that *V. vulnificus* relies on additional, incompletely understood physiological processes to adapt to host-associated environments. These findings underscore the value of genome-wide screening approaches for uncovering fitness determinants that are not readily predicted from existing virulence models.

Phenotypic analyses revealed that disruptions in motility and chemotaxis strongly impair bacterial migration (Fig. 1), confirming that these traits are tightly linked to fitness in soft tissues (Fig. 3). Although *luxO* and *barA* mutants retained normal swimming motility in vitro (Fig. 1), both mutants exhibited severe defects in persistence and dissemination within soft tissues (Fig. 3), resembling the phenotype of the motility-deficient *motX* mutant. This discrepancy indicates that regulatory pathways controlled by LuxO and BarA contribute to fitness in soft tissues through mechanisms that are independent of flagellar motility. Rather than affecting bacterial movement per se, these regulators likely coordinate additional physiological processes required for growth, persistence, or spatial expansion within host tissues.

Alterations in colony opacity among selected STM mutants further suggest that regulatory and metabolic pathways influence bacterial surface architecture. The glycosyltransferase mutant (VV1_0778::Tn) formed translucent colonies, which suggests reduced capsular polysaccharide production (Fig. 2), yet it retained the ability to disseminate within soft tissues as efficiently as WT (Fig. 3). In contrast, the *barA*::Tn mutant also exhibited a translucent colony phenotype ( Fig. 2), but IVIS analysis revealed marked defects in tissue dissemination (Fig. 3), despite this mutant retaining resistance to phagocytic stress (Fig. 4). These observations indicate that neither colony opacity nor resistance to phagocytic stress alone can reliably predict bacterial fitness in soft tissues, suggesting that additional factors independent of capsule production and motility contribute to successful growth and spread in this environment.

The IVIS analysis further revealed distinct phenotypic classes among STM-derived mutants, reflecting differential defects in persistence, dissemination, and invasion into deeper tissue compartments (Fig. 3) (12). In particular, regulatory mutants (*luxO*::Tn and *barA*::Tn) exhibited dissemination defects comparable to those of motility-deficient strains, despite differing phenotypes in other assays. LuxO is a central regulator of quorum-sensing pathways in *V. vulnificus* and has been implicated in the control of virulence-associated gene expression, including factors linked to RTX toxin regulation (18–21). Although RTX genes were not directly identified in the STM screen, impaired regulation of toxin expression or secretion may contribute indirectly to the reduced fitness of *luxO* mutants in soft tissues.

Consistent with the IVIS analysis, several STM-derived mutants displayed increased susceptibility to phagocytic stress in an HL-60-derived neutrophil model (Fig. 4). Reduced survival among motility-, regulatory-, and metabolism-defective mutants indicates that these pathways contribute to tolerance of host-derived biotic pressures encountered during early infection. Notably, *barA*::Tn retained resistance to phagocytic stress despite exhibiting impaired dissemination in soft tissues, further supporting the notion that fitness in this environment reflects the integration of multiple physiological traits rather than reliance on a single immune evasion mechanism.

GpsK is a glucosamine-specific kinase that catalyzes the conversion of glucosamine to glucosamine-6-phosphate, representing the initial step of amino sugar utilization in *Vibrio* species (23). This pathway constitutes a central component of chitin-derived nutrient metabolism and has been implicated in environmental adaptation of marine *Vibrios* (24). Although disruption of *gpsK* did not result in detectable growth impairment under *in vitro* conditions, loss of this enzyme may restrict metabolic flexibility in host-associated environments, where accessible carbon and nitrogen sources are limited. Such constraints could compromise bacterial fitness in soft tissues, contributing to the delayed tissue invasion and increased susceptibility to phagocytic stress observed for the *gpsK*::Tn mutant. These findings suggest that *gpsK* supports in vivo fitness not by enhancing basal growth capacity but by enabling metabolic adaptation to nutrient-limited and stress-rich host environments.

Taken together, the results of this study indicate that *Vibrio vulnificus* fitness in host-associated environments is governed by a network of physiological functions encompassing motility, regulatory signaling, metabolism, and stress tolerance.

Many of the identified determinants do not correspond to classical virulence factors but instead contribute to bacterial adaptation under host-imposed constraints. By linking STM-based selection with phenotypic outcomes, this work provides further insight into how environmental adaptation underpins bacterial persistence during early stages of infection and highlights the value of phenotype-centered approaches for dissecting in vivo fitness beyond canonical virulence paradigms.

## Acknowledgements

The authors thank ChatGPT (OpenAI) and the Nature Research Editing Service for assistance with English language editing and manuscript polishing. These tools were used solely to improve clarity and readability of the text and did not influence the study design, data analysis, or interpretation of results.

This work was supported by Grants-in-Aid for Scientific Research (KAKENHI) from the Japan Society for the Promotion of Science (JSPS), Grant Numbers 19K15979 and 22K14998 awarded to Kohei Yamazaki and 18H02350 awarded to Takashige Kashimoto.

## Reference

1. Baker-Austin C, Oliver JD, Alam M, et al. 2018. *Vibrio* spp. infections. Nat Rev Dis Primers 4:8. 10.1038/s41572-018-0005-8

2. Baker-Austin C, Trinanes JA, Taylor NGH, et al. 2013. Emerging *Vibrio* risk at high latitudes in response to ocean warming. Nat Clim Chang 3:73–77. 10.1038/nclimate1628

3. Stevens DL, Bryant AE. 2017. Necrotizing soft-tissue infections. N Engl J Med 377:2253–2265. 10.1056/NEJMra1600673

4. Skrede S, Bruun T, Rath E, Oppegaard O. 2020. Microbiological etiology of necrotizing soft tissue infections. Adv Exp Med Biol 1294:53–71. 10.1007/978-3-030-57616-5_5

5. Peetermans M, de Prost N, Eckmann C, et al. 2020. Necrotizing skin and soft-tissue infections in the intensive care unit. Clin Microbiol Infect 26:902–909. 10.1016/j.cmi.2020.02.005

6. Diaz JH. 2014. Skin and soft tissue infections following marine injuries and exposures in travelers. J Travel Med 21:207–213. 10.1111/jtm.12115

7. Finkelstein R, Oren I. 2011. Soft tissue infections caused by marine bacterial pathogens: epidemiology, diagnosis, and management. Curr Infect Dis Rep 13:470–477. 10.1007/s11908-011-0199-3

8. Oliver JD. 2005. Wound infections caused by *Vibrio vulnificus* and other marine bacteria. Epidemiol Infect 133:383–391. 10.1017/S0950268805003894

9. Lo HR, Lin JH, Chen YH, Chen CL, Shao CP, Hor LI. 2011. RTX toxin enhances the survival of *Vibrio vulnificus* during infection by protecting the organism from phagocytosis. J Infect Dis 203:1866–1874. 10.1093/infdis/jir122

10. Kim YR, Lee SE, Hyun K, Yeom JA, Na HS, Kim SY, et al. 2008. *Vibrio vulnificus* RTX toxin kills host cells only after contact of the bacteria with host cells. Cell Microbiol 10:848–862. 10.1111/j.1462-5822.2007.01088.x

11. Pettis GS, Mukerji AS. 2020. Structure, function, and regulation of the essential virulence factor capsular polysaccharide of *Vibrio vulnificus*. Int J Mol Sci 21:3259. 10.3390/ijms21093259

12. Yamazaki K, Kashimoto T, Kado T, Akeda Y, Yoshioka K, Kodama T. 2020. Chemotactic invasion in deep soft tissue by *Vibrio vulnificus* is essential for the progression of necrotic lesions. Virulence 11:839–847. 10.1080/21505594.2020.1782707

13. Yamazaki K, Kashimoto T, Morita M, Kado T, Matsuda K, Yamasaki M, et al. 2019. Identification of in vivo essential genes of *Vibrio vulnificus* for establishment of wound infection by signature-tagged mutagenesis. Front Microbiol 10:123. 10.3389/fmicb.2019.00123

14. Yamamoto M, Kashimoto T, Tong P, Xiao J, Sugiyama M, Inoue M, et al. 2015. Signature-tagged mutagenesis of *Vibrio vulnificus*. J Vet Med Sci 77:823–828. 10.1292/jvms.14-0655

15. Kashimoto T, Yamazaki K, Kado T, Matsuda K, Ueno S. 2021. MukB is a gene necessary for rapid proliferation of *Vibrio vulnificus* in the systemic circulation but not at the local infection site in the mouse wound infection model. Microorganisms 9:934. 10.3390/microorganisms9050934

16. Kado T, Kashimoto T, Yamazaki K, Matsuda K, Ueno S. 2019. Accurate prediction of anti-phagocytic activity of *Vibrio vulnificus* by measurement of bacterial adherence to hydrocarbons. APMIS 127:80–86. 10.1111/apm.12910

17. Yamazaki K, Kashimoto T, Kado T, Yoshioka K, Ueno S. 2022. Increased vascular permeability due to spread and invasion of *Vibrio vulnificus* in the wound infection exacerbates potentially fatal necrotizing disease. Front Microbiol 13:849600. 10.3389/fmicb.2022.849600

18. Kim SY, Lee SE, Kim YR, Kim CM, Ryu PY, Choy HE, et al. 2003. Regulation of *Vibrio vulnificus* virulence by the LuxS quorum-sensing system. Mol Microbiol 48:1647–1664.

19. Shao CP, Lo HR, Lin JH, Hor LI. 2011. Regulation of cytotoxicity by quorum-sensing signaling in *Vibrio vulnificus* is mediated by SmcR, a repressor of *hlyU*. J Bacteriol 193:2557–2565.

20. Gauthier JD, Jones MK, Thiaville P, Joseph JL, Swain RA, Krediet CJ, et al. 2010. Role of GacA in virulence of *Vibrio vulnificus*. Microbiology 156:3722–3733.

21. Choi G, Choi SH. 2022. Complex regulatory networks of virulence factors in *Vibrio vulnificus*. Trends Microbiol 30:1109–1121. 10.1016/j.tim.2022.05.009

22. Hammer BK, Bassler BL. 2003. Quorum sensing controls biofilm formation in *Vibrio cholerae*. Mol Microbiol 50:101–114.

23. Park JK, Wang LX, Roseman S. 2002. Isolation of a glucosamine-specific kinase, a unique enzyme of *Vibrio cholerae*. J Biol Chem 277:15573–15578.

24. Ran L, Wang X, He X, Guo R, Wu Y, Zhang P, Zhang XH. 2023. Genomic analysis and chitinase characterization of *Vibrio harveyi* WXL538: insight into its adaptation to the marine environment. Front Microbiol 14:1121720. 10.3389/fmicb.2023.1121720

